# Cold shock induces a terminal investment reproductive response in *C. elegans*

**DOI:** 10.1101/2021.09.11.459896

**Authors:** Leah Gulyas, Jennifer R. Powell

## Abstract

Challenges from environmental stressors have a profound impact on many life-history traits of an organism, including reproductive strategy. Examples across multiple taxa have demonstrated that maternal reproductive investment resulting from stress can improve offspring survival; a form of matricidal provisioning when death appears imminent is known as terminal investment. Here we report a reproductive response in the nematode *Caenorhabditis elegans* upon exposure to acute cold shock at 2°C, whereby vitellogenic lipid movement from the soma to the germline appears to be massively upregulated at the expense of parental survival. This response is dependent on functional TAX-2;TAX-4 cGMP-gated channels that are part of canonical thermosensory mechanisms in worms and can be prevented in the presence of activated SKN-1/Nrf2, the master stress regulator. Increased maternal provisioning promotes improved embryonic cold shock survival, which is notably suppressed in animals with impaired vitellogenesis. These findings suggest that cold shock in *C. elegans* triggers terminal investment to promote progeny fitness at the expense of parental survival and may serve as a tractable model for future studies of stress-induced progeny plasticity.

## Introduction

Environmental stressors can severely jeopardize the ability of an organism and its progeny to survive and reproduce, endangering organismal fitness. Consequently, organisms must launch effective stress response mechanisms to mitigate damage, promote recovery, and/or ensure progeny survival. A traditional view of life-history trade-offs assumes that challenge from a stressor should invoke a response in which resources are shuttled away from reproduction to support organismal defense and repair [1]. While this approach temporarily decreases reproductive fitness, improved long-term survival may allow for greater reproduction post-recovery. However, in a stressful environment where survival is perceived to be unlikely, an alternate strategy may instead be to funnel resources into reproduction, ensuring maximal output prior to parental death and subsequent survival of the F1 generation [1].

The benefit of this strategy has been demonstrated in a number of organisms in different environmental conditions [2]. Both biotic (i.e. pathogen exposure) and abiotic (thermal, osmotic) stress conditions may potentiate molecular changes to reproduction. Alterations ranging from epigenetic modifications to simple maternal cytosolic investments of macromolecules and nucleic acids have been associated with increased offspring resilience to ensuing threats [2,3]. Yet despite myriad examples, the precise mechanisms by which stressors induce these responses are still widely under exploration.

The nematode *Caenhorhabditis elegans* recently arose as an ideal model for investigating stress-induced parental progeny investment. Several studies have detailed molecular tactics by which worms modify and provision gametes and embryos in response to dietary restriction, infection by pathogenic pseudomonads and microsporidia, and osmotic stress [4–7]. Among an array of environmentally-relevant stressors, though, the reproductive impacts of thermal stress (specifically cold stress) on *C. elegans* have not been closely assessed. Both flies and mice employ reproductive strategies to respond to cold, with flies inducing a more general stress response of reproductive dormancy and mice depositing epigenetic modifications in sperm to promote retention of brown adipose tissue favoring increased metabolic rates in offspring [8,9]. Given the likelihood of *C. elegans* regularly encountering diurnal and seasonal temperature fluctuations, it seems probable that worms may also have evolved rigorous reproductive programs to deal with cold exposure.

Several aspects of cold response in *C. elegans* have been well-characterized in the literature, including thermotaxis and habituation [10–12]. Neurons and other tissues rely on a multitude of signaling mechanisms for these physiological responses to occur, which are in turn supported by a bevy of channels and other proteins. These processes are described in detail in a recent review from Takeishi et al. [12]. Only lately, though, have links between cold and reproduction begun to emerge. Sonoda et al. [13] demonstrate that the presence and integrity of sperm is required for normal cold tolerance by exerting an effect through multiple tissues, including ASJ neuronal activity. At the level of population dynamics, it has also been reported that *C. elegans* warming from a 2-hour cold exposure enter a state of programmed death which is thought to result in kin selection for the survival of younger (and presumably more reproductively fit) animals [14]. However, the extent to which acute cold stress directly impacts reproductive capacity in these animals remains unknown. As alternating periods of cold and moderate temperatures preceding a winter season may comprise a substantial portion of the short reproductive window in *C. elegans*, this necessitates further study.

We have previously shown that acute cold shock at 2°C causes loss of intestinal pigmentation, immobility, and reproductive disruption in wild-type hermaphrodite worms. In many cases, this phenotypic program results in lethality [15]. Here we extend this work by characterizing the induction of these phenotypes via the neuronal TAX-2/TAX-4 thermotransduction channel, mediated by the canonical stress response regulator SKN-1/Nrf2. We further describe a role for the normal vitellogenesis machinery in reallocating pigmented intestinal lipid supplies to the germline, which appears to come at the expense of parental survival following cold exposure. Overall, our findings suggest that acute cold shock provokes a terminal investment reproductive response in *C. elegans*, making this discovery an exciting opportunity to explore the phenomenon in a genetically tractable system.

## Results

### Acute cold shock causes drastic phenotypic alterations

The duration of cold exposure for young adult hermaphrodite *C. elegans* at 2°C is negatively correlated to post-shock survival rates [15]. Wild-type hermaphrodite worms exposed to a 4-hour cold shock (CS) do not initially display high mortality rates (**Fig. 1a**); this allows observation of a range of phenotypic transitions as they recover from the limited-duration cold stress at their preferred temperature of 20°C. One of the most striking phenotypes exhibited in post-cold shock (post-CS) animals during the recovery period is a dramatic decrease in pigmentation in the normally highly pigmented intestine, so that the body becomes almost entirely clear (**Fig. 1b, c**) [15]. This is often accompanied by motor and reproductive disruptions such as mobility loss, withering of the gonad arms, decreased number of internal embryos, and the eventual death of about 30% of the population (**Fig. 1a-d**) [15]. It should be noted that these phenotypic responses do not appear to be due to any relative heat shock following the transition from 2°C to 20°C as the expression of GFP-tagged HSP-4 (heat shock protein) is not induced following cold shock (**Fig. 1e**). Interestingly, some CS wild-type animals regain pigmentation after clearing; these worms do not die and display a general reversal of the other negative impacts of cold shock (**Fig. 1b**)[15]. We sought to better understand the factors regulating the post-CS recovery program in wild-type worms, focusing particularly on the functional role of pigmentation loss and the genetic components involved in producing it.

**Figure 1.**
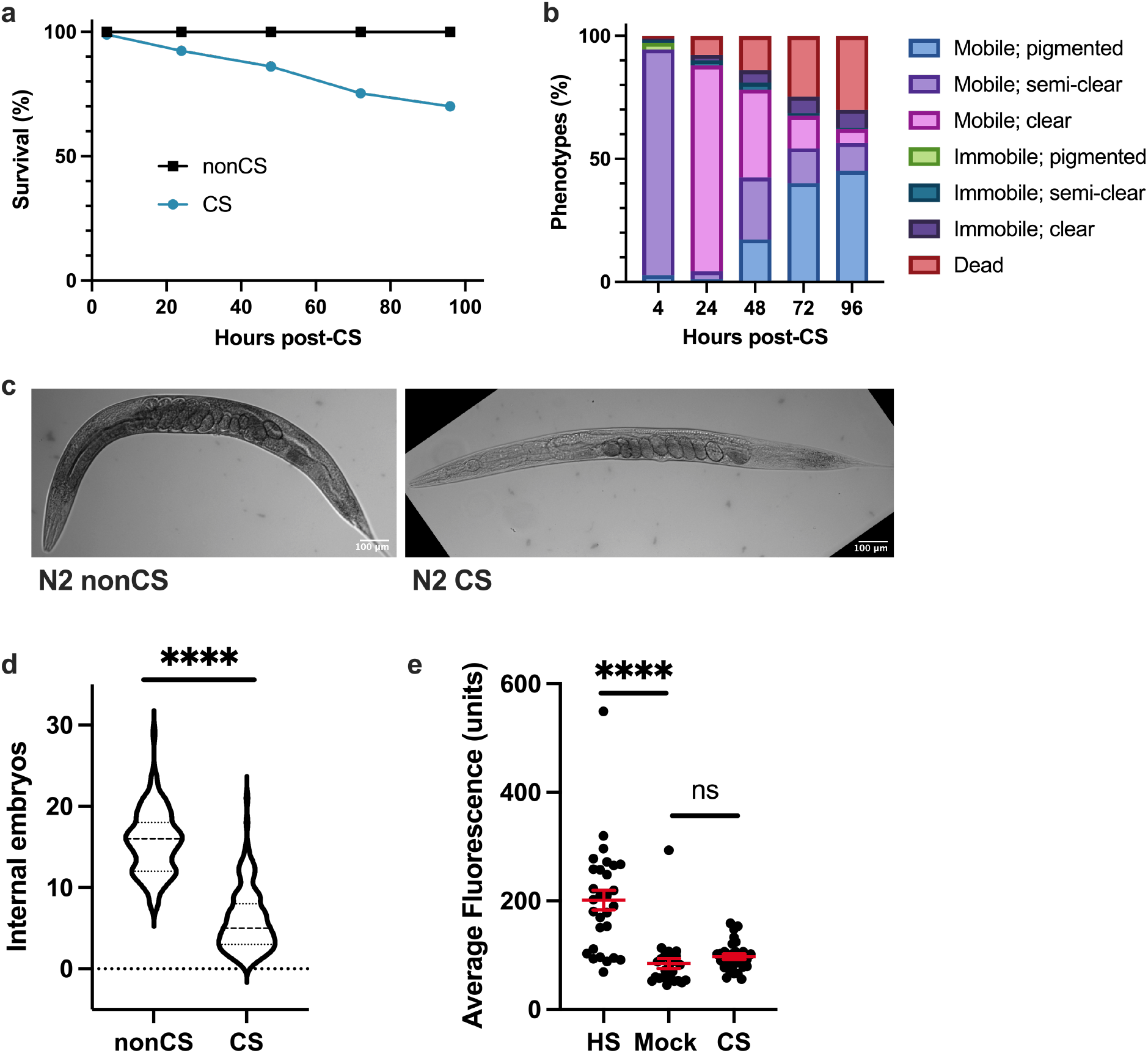
Cold-shocked worms show decrease in survival and characteristic phenotypic alterations. N2 young adult hermaphrodites were shifted from 20°C to 2°C for a 4 h cold shock (CS) and thereafter recovered at 20°C for 96 h with assessment of **(a)** survival and **(b)** phenotypic alterations (n = 177). Death and immobility were assayed by nose tap; worms were considered to be immobile if the tap elicited slight movement in the head region but no other body movement, and dead worms were completely unresponsive. **(c)** Representative images of young adult N2 hermaphrodites following cold shock. **(d)** Average number of internal embryos per worm in CS versus nonCS conditions (Mann-Whitney U test: U=604, two-tailed P < 0.0001). **(e)** Induction of heat shock protein HSP-4::GFP 12 hours after either 2 h heat shock at 35°C, 4 h cold shock at 2°C, or no treatment (n ≥ 26 per condition; Kruskal-Wallis test: H=33.86, P < 0.0001; Dunn’s Multiple Comparison Test: ****P < 0.0001). Error bars are mean ± s.e.m.

### Cold shock induces lipid reallocation from somatic tissues to the germline

Since pigmentation in the *C. elegans* intestine is due in part to the presence of lipid-storing fat droplets [16], we wondered whether the decrease in pigmentation in cold-shocked worms corresponds to a depletion of intestinal lipid supplies, potentially as a result of increased metabolism meant to fuel post-cold shock recovery. Using Nile Red, which accumulates and fluoresces in hydrophobic environments [17], as an indicator of lipid content, we therefore analyzed the fat stores of worms 12 hours following CS (note that all cold shocks were performed for 4 h at 2°C). We indeed observed a significant decrease (P < 0.0001) in the average lipid content per worm (**Fig. 2a**); visually, this presents as an overall reduction in average fluorescence that is most striking in the intestine (**Fig. 2c**). However, we also unexpectedly noted that the lipid content of the gonad appears to concomitantly increase, marked by the presence of fluorescent (and therefore lipid-rich) internal embryos in the gonads of many cold-shocked worms that were mostly absent in their non-cold shocked counterparts (**Fig. 2b, c**). These embryos are somewhat sporadic, usually accounting only for a proportion of all embryos in the germline and are interspersed with non-fluorescent embryos. Importantly, the percentage of fluorescent internal embryos per worm was found to be elevated two-fold in CS worms relative to that of nonCS controls (**Fig. 2b**). These observations suggest that rather than just metabolically depleting lipid supplies, *C. elegans* may also reallocate lipids from the intestine to the germline. To further analyze this process spatiotemporally, we performed a time course of Nile Red staining following cold shock (**Fig. 2c**). Upon entry into 2°C shock, young adult worms enter a “chill coma,” ceasing virtually all movement [15,18]. Following exit from cold shock, most worms gradually resume movement over the first 30 minutes of recovery at 20°C. At this point, the distribution of Nile Red-staining lipids is indistinguishable from non-cold shocked controls, indicating that lipid reallocation occurs after, rather than during, the cold shock. By 4 hours post-cold shock, we observed visually decreased but rarely absent fluorescence in the intestine; at this time, many worms have brightly fluorescing proximal oocytes, suggesting that lipids are beginning to relocate from the intestine into the germline. This process of reallocation appears to continue over the next 14+ hours, with many 12 h-recovered worms containing fluorescent embryos and oocytes but only slight intestinal fluorescence, and most 18 h-recovered worms fluorescing almost exclusively in the embryos. Though it is unclear what other factors (e.g. metabolism) may contribute to the overall pigmentation loss, these observations suggest that the dramatic decrease in intestinal pigmentation results in large part from the reallocation of lipids from somatic tissues to the germline during the recovery phase following a severe cold shock.

**Figure 2.**
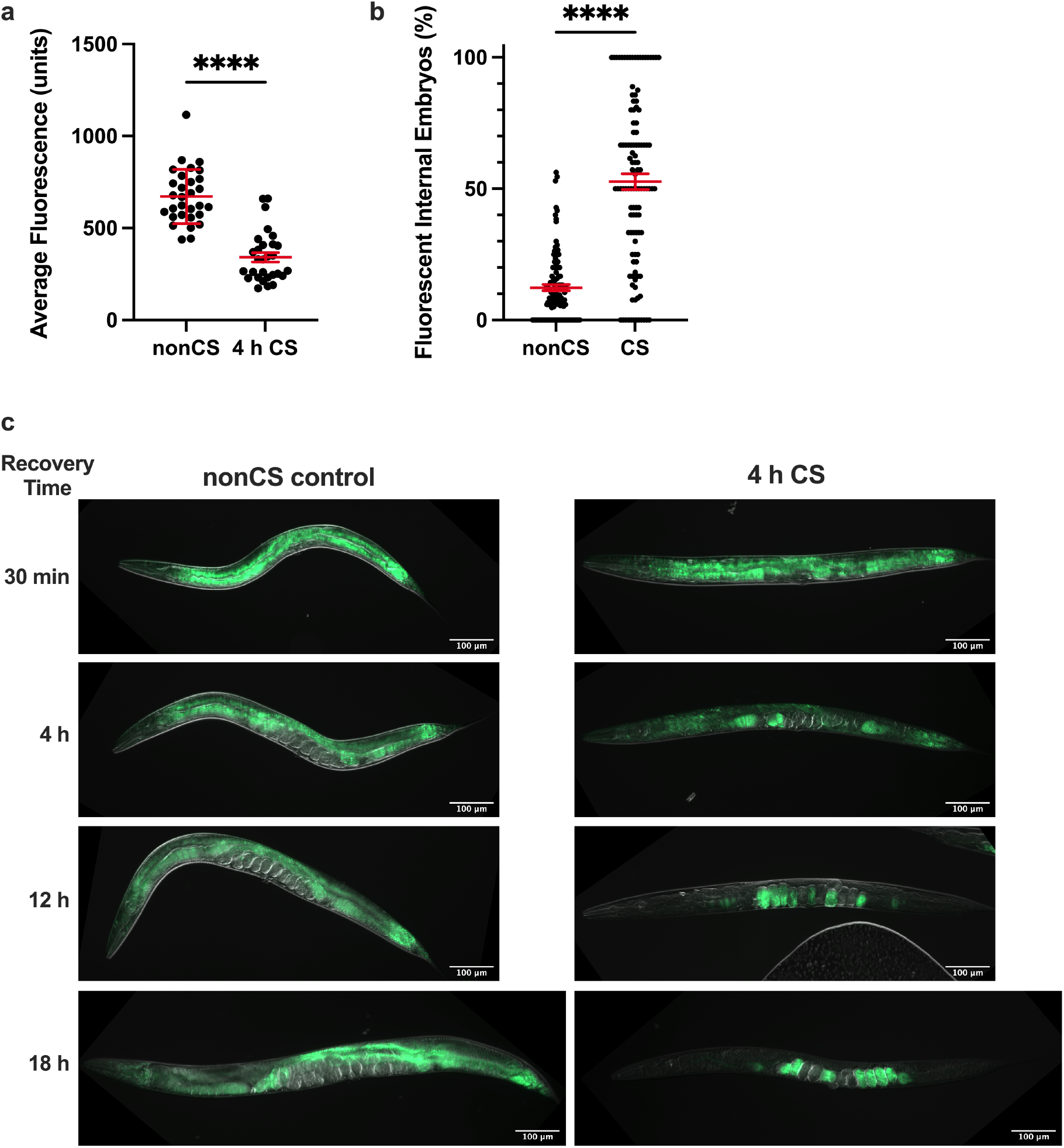
Cold shock recovery is associated with lipid localization shifts from the intestine to the germline. Young adult N2 hermaphrodites were exposed to 2°C cold shock or control nonCS conditions (20°C) for 4 h and recovered before Nile Red staining and imaging for lipid content. **(a)** Average fluorescence per worm (n ≥ 29 per condition; Mann-Whitney U test: U=48; two-tailed P < 0.0001) and **(b)** percent of internal fluorescent embryos per worm (n ≥ 111; Mann-Whitney U test: U=1885; two-tailed P < 0.0001) were quantified 12 h post-CS. Error bars are mean ± s.e.m. **(c)** Representative images of Nile Red-stained worms at indicated time points post-CS show progression of intestinal lipid loss and relocalization to the embryos.

### Thermosensation is required to initiate the cold shock response

We next asked how wild-type *C. elegans* perceive and interpret cold shock as a signal for lipid reallocation. The cGMP-gated channel subunits TAX-2 and TAX-4 are important for thermosensation and acquired cold tolerance in worms, acting specifically in the ASJ neurons [11,19,20]. We reasoned that worms might also rely on a neuronal mechanism of sensation that acts through the TAX-2/TAX-4 channel to induce the cold shock response in the absence of prior cold exposure. *tax-2*(*p671*); *tax-4*(*p678*) double loss-of-function mutants were therefore tested for sensitivity to cold shock. Not only do these mutants show 100% survival following cold shock (**Fig. 3a**), but, unlike wild-type worms, they also do not display any significant difference (P > 0.9999) in overall lipid content 12 h after CS, nor the striking lipid reallocation phenotype characterized by large numbers of fluorescent embryos (**Fig. 3b-d**). Interestingly, despite this resilience to shock, *tax-2*(*p671*); *tax-4*(*p678*) mutants still show decreased internal embryo quantities, though not to the same degree as N2 worms (**Fig. 3e**), suggesting that cold exposure has some additional impact on fertility independent of this thermosensation pathway. Taken together, though, these data support the hypothesis that fat reallocation as a stress response following CS may depend on a neuronal mechanism for induction. Since TAX-2/TAX-4 plays a role in temperature sensation, we speculate that canonical cold thermosensation is required for lipid reallocation to take place.

**Figure 3.**
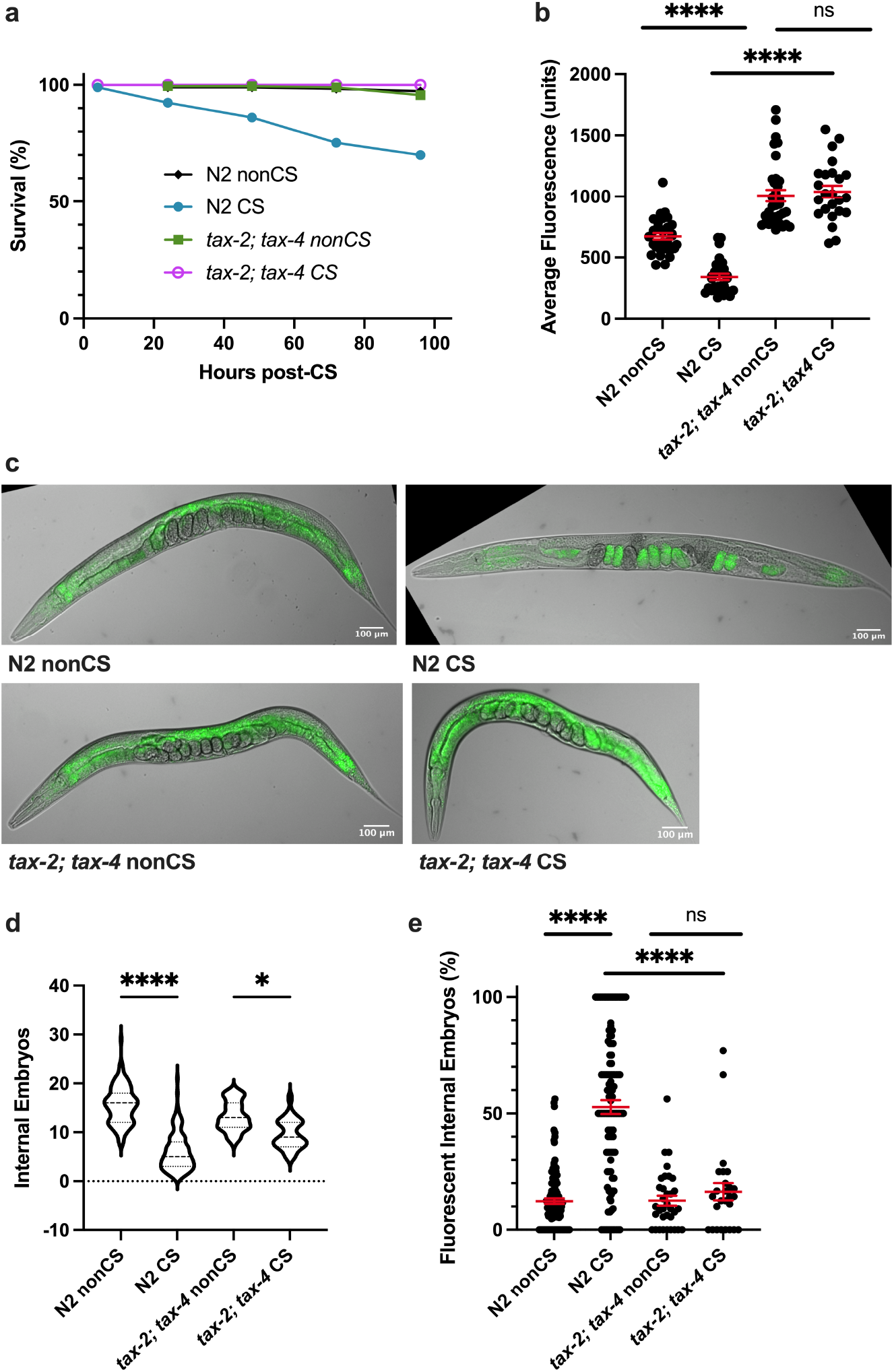
TAX-2; TAX-4-mediated thermosensation is required for lipid relocalization following cold shock. **(a)** *tax-2*(*p671*); *tax-4*(*p678*) double loss-of-function mutants were cold shocked and recovered while monitoring survival rates (n ≥ 120 worms per condition). **(b)** At 12 h post-CS or nonCS control, *tax-2*(*p671*); *tax-4*(*p678*) were Nile Red lipid-stained and the average fluorescence per worm was quantified (Kruskal-Wallis test: H = 83.83, P < 0.0001; Dunn’s Multiple Comparison test: ****P < 0.0001). **(c)** Representative images of Nile Red staining show retention of fluorescence in the intestines of *tax-2*; *tax-4* mutants. **(d)** Number of embryos (Kruskal-Wallis test: H = 171.1, P < 0.0001; Dunn’s Multiple Comparison test: *P = 0.0484, ****P < 0.0001) and **(e)** percent fluorescent internal embryos (Kruskal-Wallis test: H = 100.1, P < 0.0001; Dunn’s Multiple Comparison test: ****P < 0.0001) were quantified per worm from Nile Red images (n ≥ 24 worms per condition for b-e; error bars are mean ± s.e.m.).

### SKN-1 promotes cold stress resistance

After determining a requirement for neuronal signaling in cold-shock-induced lipid reallocation, we next wondered what intermediate factors were needed to translate the cold stimulus into a signal to mobilize resources. The master regulatory transcription factor SKN-1/Nrf2 coordinates a return to homeostasis following a variety of stresses, including oxidative stress [4], pathogen infection [21,22], and osmotic stress [23]. We therefore predicted that *skn-1* would perform a similar function during cold stress recovery. Indeed, *skn-1*(*lax188*) gain-of-function mutants [24] are highly resistant to cold shock, with nearly 100% survival rates 96h post-CS (**Fig. 4a**). Consistent with a retention of overall lipid stores by Nile Red Staining (**Fig. 4b,d**), these mutants did not contain significantly greater numbers of fluorescing internal embryos following cold shock (P > 0.9999), though they did have a marginal reduction in the number of internal embryos (**Fig. 4d-f**). Conversely, *skn-1*(*mg570*) loss-of-function mutants displayed a wild-type cold shock response, with significant reallocation (P < 0.00001) of lipids to the germline and loss of somatic fats (**Fig. 4c-f**). These data suggest that the *skn-1a* isoform, which is knocked out by the *mg570* mutation [25], is not required for lipid mobilization in response to cold shock. However, SKN-1 activation protects cold shocked worms from recovery phase lethality and prevents lipid reallocation.

**Figure 4.**
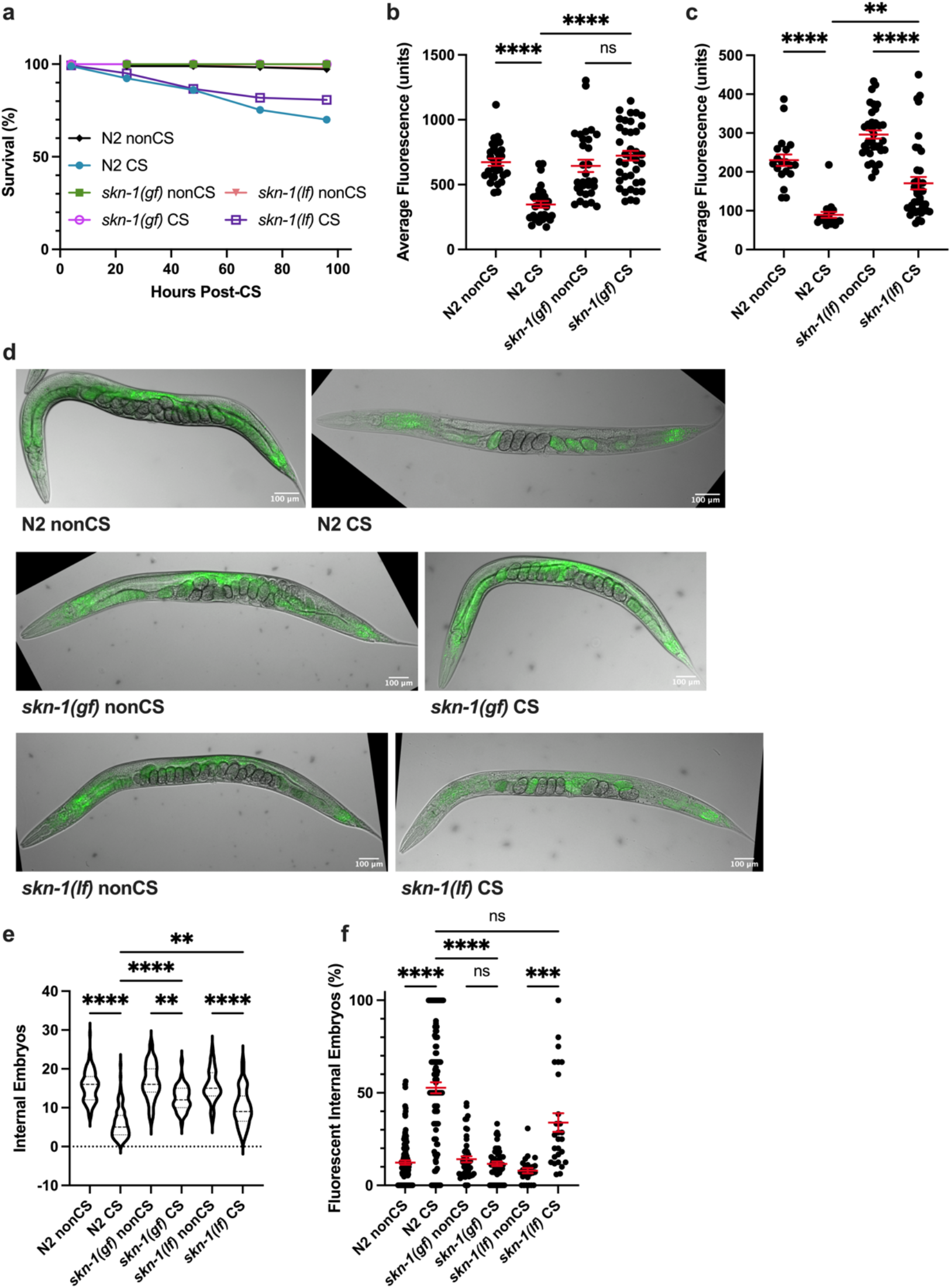
Activated SKN-1 prevents lipid relocalization following cold shock. **(a)** *skn-1*(*lax188*) gain-of-function mutants and *skn-1*(*mg570*) loss-of-function mutants were cold shocked and recovered while monitoring survival rates (n≥120 worms per condition). **(b)** At 12 h post-CS or control nonCS, *skn-1*(*lax188*) and *skn-1*(*mg570*) were Nile Red lipid-stained and average fluorescence per worm quantified (N2 v. *skn-1*(*lax188*)-Kruskal-Wallis test: H = 47.46, P < 0.0001; Dunn’s Multiple Comparison test: ****P < 0.0001; N2 v. *skn-1*(*mg570*)-Kruskal-Wallis test: H=64.55, P < 0.0001; Dunn’s Multiple Comparison test: **P = 0.0064, ****P < 0.0001). **(c)** Relative to wild-type N2s, representative images of *skn-1*(*lax188*) mutants show maintenance of intestinal fluorescence contrary to *skn-1*(*mg570*), which show lipid relocalization. **(d)** Number of embryos (Kruskal-Wallis test: H = 223.1, P <0.0001; Dunn’s Multiple Comparison test: \*\**P_skn-1(gf) nonCS vs. CS_* = 0.0020, \*\**P_N2 CS vs. skn-1(lf) CS_* = 0.0014, **** P < 0.0001) and **(e)** percent of fluorescent internal embryos (Kruskal-Wallis test: H = 132.0, P < 0.0001; Dunn’s Multiple Comparison test: ***P = 0.0002, ****P < 0.0001) were quantified per worm from Nile Red images (n ≥ 26 worms per condition for b-e; error bars are mean ± s.e.m.).

### Lipid reallocation results from upregulated vitellogenesis following cold shock

The process governing the normal movement of lipids from the soma to the germline is vitellogenesis, requiring the vitellogenin proteins coded for by *vit-1-6* genes. Once coupled to somatic lipid supplies, the resulting lipoprotein complexes shuttle to the germline [26]. We hypothesized that the lipid mobilization induced by cold shock is a result of the normal vitellogenesis machinery being commandeered as part of a stress response.

One of the major vitellogenins is VIT-2, which generates a 170 kD yolk protein product known as YP170B [27]. We predicted that loss of *vit-2* would reduce lipid reallocation following CS. Since preventing reallocation in our previous mutants also increased post-CS survival, we expected that *vit-2* loss of function would enhance protection from CS. Consistent with these hypotheses, impairment of vitellogenesis in *vit-2*(*ok3211*) loss-of-function mutants produced phenotypes similar to *tax-2*(*p671*); *tax-4*(*p678*) animals. In addition to maintaining high survival rates, CS *vit-2*(*ok3211*) mutants did not undergo a substantial loss of intestinal lipid supplies, nor did they exhibit a marked increase in internal embryo fluorescence (despite a modest decrease in internal embryo counts) (**Fig 5a-e**).

**Figure 5.**
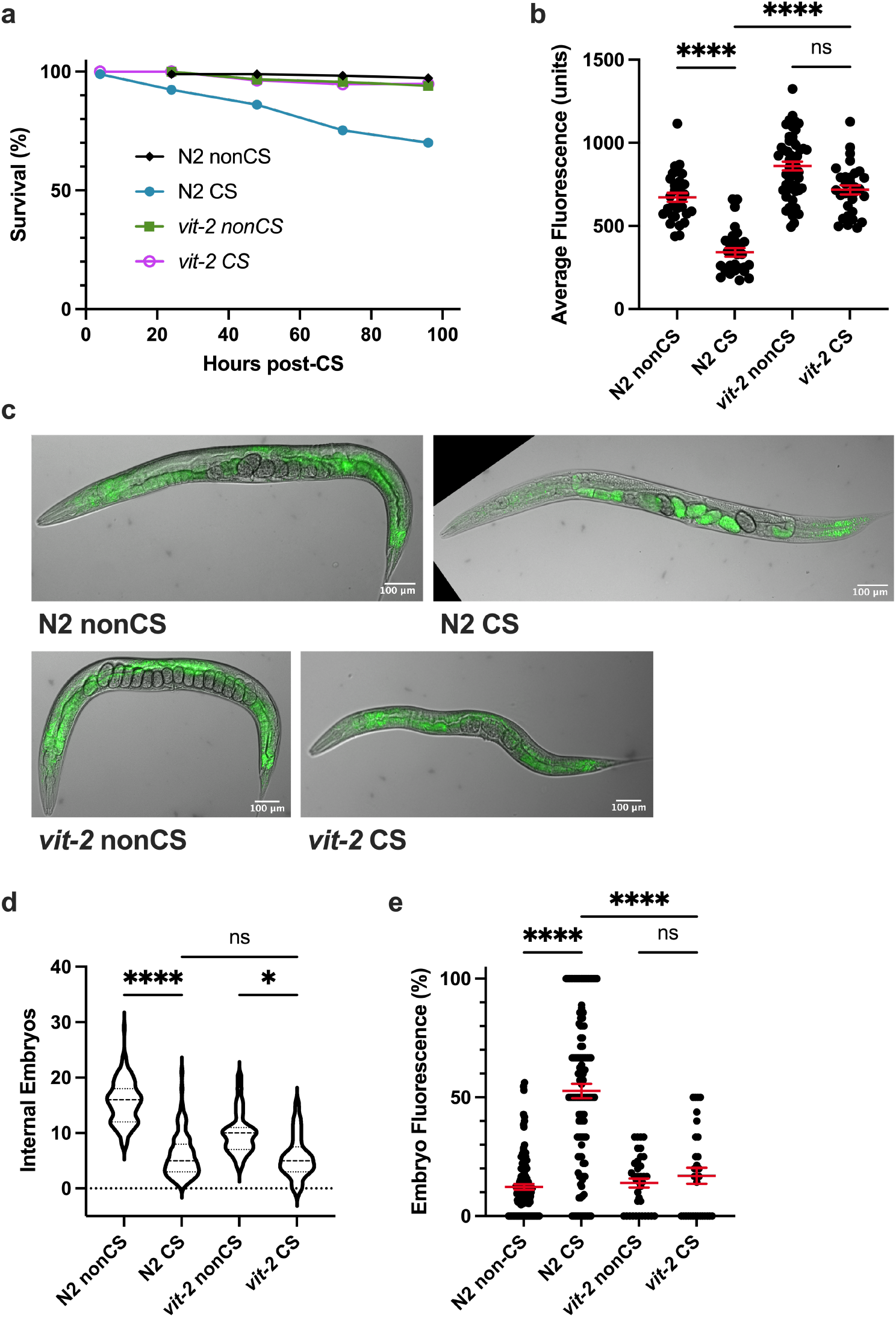
VIT-2-regulated vitellogenesis during recovery promotes lipid relocalization and impairs survival. **(a)** *vit-2*(*ok3211*) loss-of-function mutants were cold shocked and recovered while monitoring survival rates (n ≥ 130 worms per condition). **(b)** At 12 h post-CS or nonCS, *vit-2*(*ok3211*) were Nile Red lipid-stained and average fluorescence per worm quantified (n ≥ 31 worms per condition; Kruskal-Wallis test: H = 75.18, P < 0.0001; Dunn’s Multiple Comparison test: ****P<0.0001). Error bars are mean ± s.e.m. **(c)** Images of Nile Red Staining show intestinal lipid retention in *vit-2* mutants. **(d)** Number of embryos (Kruskal-Wallis test: H = 169.9, P < 0.0001; Dunn’s Multiple Comparison test: *P = 0.0149, ****P < 0.0001) and **(e)** percent of fluorescent internal embryos (Kruskal-Wallis test: H = 101.5, P < 0.0001; Dunn’s Multiple Comparison test: ****P < 0.0001) were quantified per worm from Nile Red images (n ≥ 31 for b-e; error bars are mean ± s.e.m.).

Since other vitellogenin transcripts produce different lipoprotein forms for movement to the germline, we decided to additionally test whether a member of the YP170A-generating class of vitellogenins would recapitulate our results with *vit-2.* To this end, we analyzed the CS phenotypes of *vit-5*(*ok3239*) loss-of-function animals. Indeed, we found that as in *vit-2* mutants, survival was rescued in these animals following CS, corresponding with a retention of somatic lipids and inhibition of embryonic lipid reallocation (**Fig. 6a-c,e**). As with other mutants that prevent lipid reallocation, there was still an effect of cold in reducing overall embryo output 12 h post-CS, hinting that temperature impacts fertility independent of the other reproductive alterations exhibited by wild-type worms (**Fig. 6d**).

**Figure 6.**
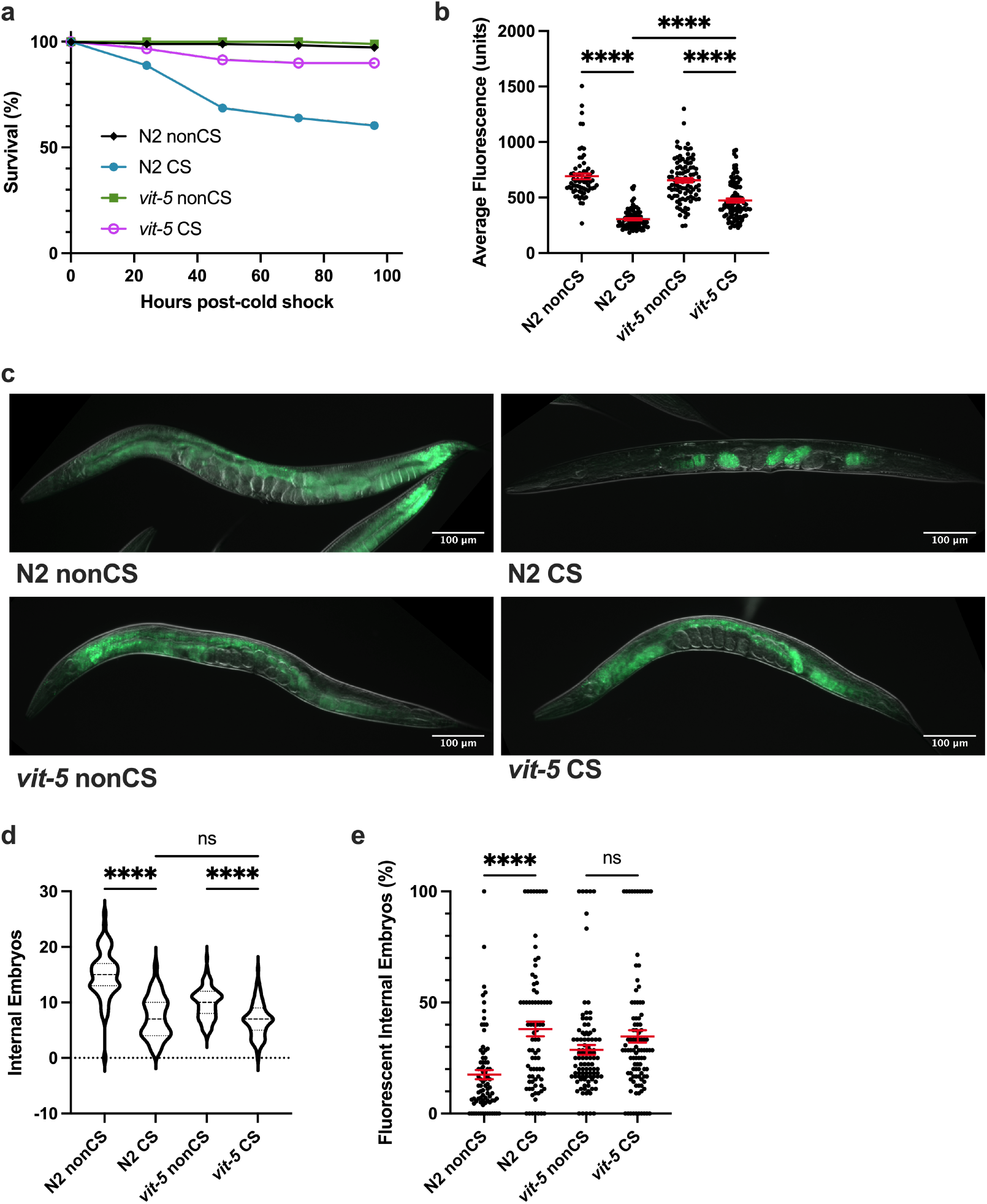
VIT-5 vitellogenin family also promotes embryo lipid reallocation during cold shock recovery. **(a)** *vit-5*(*ok3239*) loss-of-function mutants were cold shocked and recovered while monitoring survival rates (n ≥ 170 worms per condition). At 12 h post-CS or control nonCS, *vit-5*(*ok3239*) were Nile Red lipid-stained and average fluorescence per worm quantified (n ≥ 67 worms per condition; Kruskal-Wallis test: H = 172.0, P < 0.0001; Dunn’s Multiple Comparison test: *****P < 0.0001. Error bars are mean ± s.e.m. **(c)** Representative Nile Red staining shows intestinal fluorescence in *vit-5(ok3239*) following cold shock. **(d)** Number of embryos (Kruskal-Wallis test: H = 137.2, P < 0.0001; Dunn’s Multiple Comparison test: ****P < 0.0001) and **(e)** percent of fluorescent internal embryos (Kruskal-Wallis test: H = 37.58, P < 0.0001; Dunn’s Multiple Comparison test: H=37.58, ****P<0.0001) were quantified per worm from Nile Red images (n ≥ 67 worms per condition; error bars are mean ± s.e.m.).

### Cold shock response is a form of a terminal investment

Our data thus far suggest that upon recovery from acute CS, wild-type *C. elegans* induce a reproductive phenotype whereby somatic lipid supplies are massively relocalized to the germline using the normal vitellogenesis machinery. This appears to come at the expense of the parent’s own mortality, as retaining somatic lipids by preventing thermosensation or vitellogenesis is sufficient to rescue survival. Such a tradeoff between survival and reproduction is redolent of the *terminal investment hypothesis* (reviewed in Gulyas and Powell, 2019 [2]), which predicts that in some instances of acute stress where the likelihood of death is high, organisms can preferentially funnel resources to reproduction to maximize reproductive fitness at the cost of survival.

To determine whether cold stress-associated phenotypes in *C. elegans* are an example of such a process, we eliminated all potential for reproductive investment by assaying sterile *glp-1*(*e2141*) and *glp-1*(*q231*) worms to ask whether these worms still exhibited lipid loss or death upon cold shock. Strikingly, the absence of a germline in these animals completely prevented both intestinal clearing and death, again confirming that lipid movement from the soma to the germline is associated with parental lethality (**Fig. 7a**). If resource reallocation to the progeny is indeed meant to increase reproductive fitness in inclement conditions, there should be some benefit to the progeny of CS worms that offspring of nonCS parents *do not* receive. We speculated that in the case of environmental CS, a sudden, seasonal, cold to warm cycle might signal likelihood of future cold conditions that would impede the ability of embryos to hatch and survive to reproductive age. We therefore devised an assay to test for the relative fitness of embryos in cold conditions depending on whether they came from nonCS or CS parents and were thus more likely to have received lipid provisioning. To do this, embryos were collected from nonCS and CS parents within a 2 h time window when post-CS lipid reallocation seems to peak and subsequently exposed them to a cold stress of 24 hours. We then assessed hatching rates 24 hours following the embryonic cold shock. Excitingly, embryos coming from parents that had experienced cold shock prior to reproduction exhibited a small but significant increase in hatching rates relative to their counterparts from nonCS hermaphrodite parents. Furthermore, preventing lipid reallocation by impairment of vitellogenesis in either *vit-2*(*lf*) or *vit-5*(*lf*) was able to substantially ablate this effect, suggesting that the improved survival is attributable specifically to lipid investment in the F1 generation (**Fig. 7b-c**). Altogether, it appears that while preventing embryonic lipid endowment may promote adult survival post-CS, offspring that go on to experience future inclement conditions suffer diminished survival rates, underscoring the evolutionary advantage of terminal investment as a reproductive strategy.

**Figure 7.**
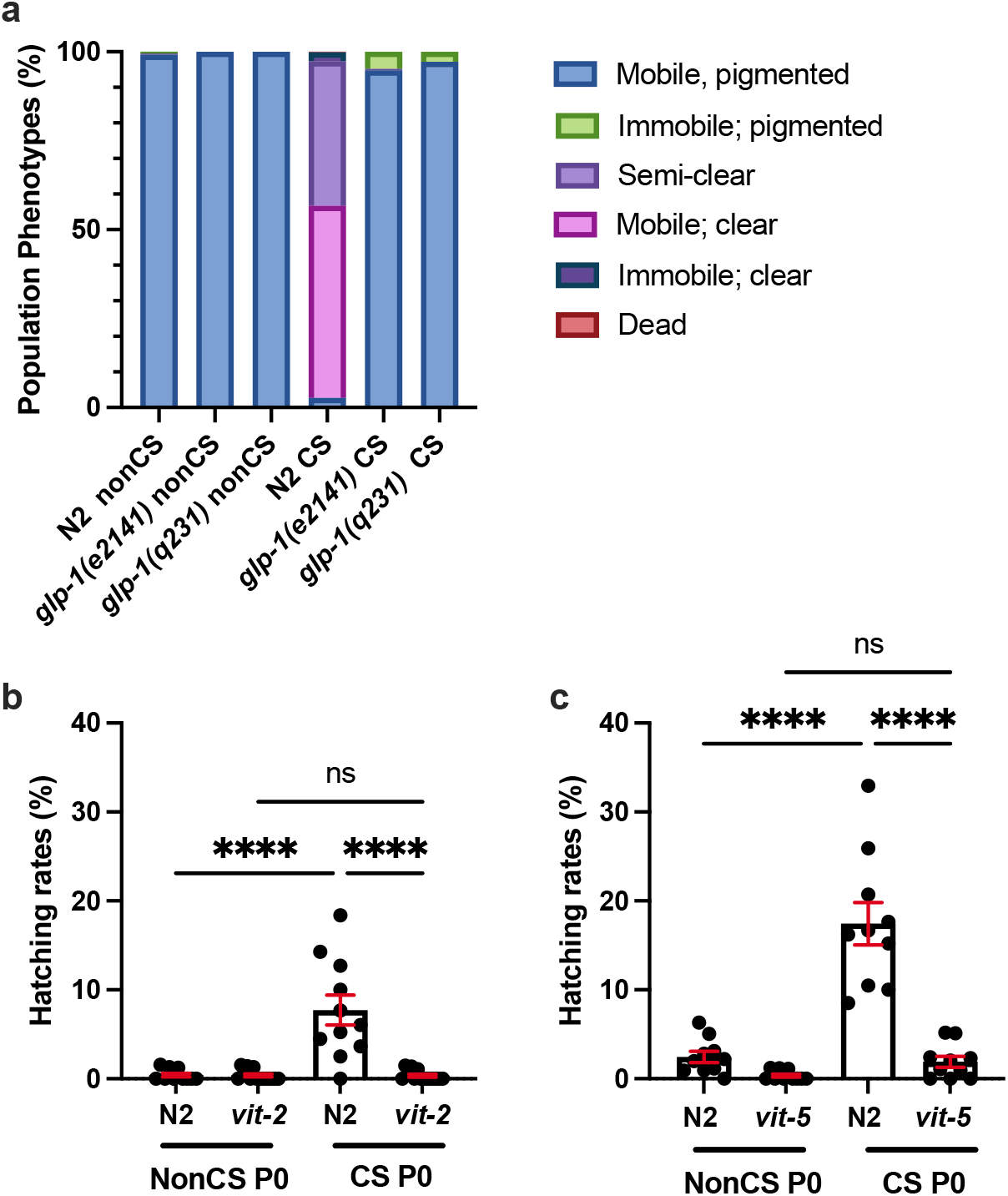
Cold shock response is a form of terminal investment. **(a)** *glp-1*(*e2141*) and *glp-1*(*q231*) were grown at the restrictive temperature (25°C) to induce sterility and then habituated at 20°C for 4 h. Worms were then cold shocked or mock shocked (nonCS) and recovered at 20°C for 24 h and phenotypes were scored (n ≥ 197 worms for all conditions). **(b)** Young adult hermaphrodite N2 and *vit-2*(*ok3211*) (P0) were cold shocked or mock shocked (nonCS) for 4 h and recovered for 16 h. Embryos from P0 treatments were collected from 16-18 h and cold shocked for 24 h. Embryos were allowed to recover 24 h at 20°C and then the number of hatched embryos was quantified. Data points represent rates as percent hatched for a plate containing 50-100 embryos (n ≥ 10 plates per condition; 2-way ANOVA (F(3,32) = 20.87, P < 0.0001) with Tukey multiple comparison test (****P < 0.0001). Bars are mean ± s.e.m. **(c)** Performed as in **b** with N2 and *vit-5*(*ok3239*) worms (n = 10 plates per condition; 2-way ANOVA (F(3,27) = 35.28, P < 0.0001) and Tukey’s multiple comparison test (****P < 0.0001).

## Discussion

### Cold shock response overlaps with other stress response phenotypes

We have provided here the first evidence that *C. elegans*, upon acute cold shock (CS), is able to improve the success of progeny survival during future cold exposure by mobilizing lipids to the germline as reproductive provisioning. This process seems to depend on neuronal sensation of the cold temperature, which permits a shift of lipid localization from the soma to the germline. Lipid reallocation is mediated by vitellogenins, and despite an overall decrease in embryo output, the embryos in these worms are more resilient to future cold exposure and display more robust hatching. The switch to a sudden last reproductive event and resulting parental mortality is consistent with the adoption of a more semelparous lifestyle [28]. In this instance, potentiation by a severe environmental stress to induce a higher quality (rather than quantity) final reproductive investment suggests that C. elegans has evolved a terminal investment response to deal with acute cold exposure.

Though, to our knowledge, no previous study has linked terminal investment to *C. elegans,* there are other documented phenomena characterized by a similar trade off. For example, the “age-dependent somatic depletion of fat (ASDF)” phenotype is characterized by age-dependent changes in lipid homeostasis that are induced during both starvation and oxidative stress and are highly reminiscent of post-cold-shock reallocation of lipids from the intestine to the germline [24]. As is also the case with cold-shock-induced terminal investment, reduction of vitellogenic transcripts (including *vit-2*) suppresses lipid mobilization. Lipid reallocation during ASDF does appear to negatively impact parental survival, but it is unclear whether this phenotype also holds functional significance for offspring survival. Surprisingly, while the process governing the ASDF phenotype and cold shock recovery are both regulated by the master stress response transcription factor SKN-1/Nrf2, activating SKN-1 has opposite effects. When measuring ASDF, *skn-1*(*gf*) stimulates the shift of lipids from the intestine to the germline [24]; in contrast, *skn-1*(*gf*) prevents lipid mobilization and promotes intestinal lipid sequestration following cold shock. It is possible that the role of SKN-1 varies with age since ASDF occurs in much older adults than those analyzed following cold stress. Although regulation may be via distinct pathways, the existence of mechanistically similar lipid homeostasis responses to starvation, oxidative stress, and cold stress suggests that terminal investment may be a more generalized response to severe stressors in *C. elegans*.

Despite the fact that terminal investment requires extremely high parental resource investment, there are other similar and less severe forms of maternal provisioning in *C. elegans*. For example, osmotic stress shifts embryonic contents to include less glycogen and more glycerol, a cryoprotectant [5]. It is particularly relevant to our study that mild nutrient deprivation in hermaphrodites leads to intergenerational plasticity in which embryos tend to be slightly larger and offspring are somewhat protected from the effects of larval starvation [4]. Remarkably, this corresponds to decreased maternal insulin signaling and upregulated vitellogenin provisioning in the germline, much as we report during cold exposure [29]. It will be interesting in future experiments to dissect how thermosensory signaling may hyperactivate vitellogenesis upon acute cold stress, especially as it relates to different vitellogenin yolk protein products. Additionally, analyzing how various stressors impact and modify embryonic composition will be important for understanding the extent to which maternal investments are conserved. Taken together, it is clear that stressors in the parental generation can have remarkable consequences on reproductive and survival tradeoffs as well as offspring physiology and plasticity.

### Terminal investment in response to cold may have evolved in regularly fluctuating environmental conditions

Since the preeminent goal of survival is reproduction, terminal investment is sensible from a fitness standpoint. But why would cold shock elicit this phenotype? Experimental evolution in *C. elegans* reveals that deterministic maternal effects (those that attempt to successfully invest offspring for survival in a particular environment) evolve in response to predictably fluctuating environments [30]. Thus, we conjecture that predictably encountered cold environments, such as those during seasonal freeze-thaw cycles in temperate regions, are likely to have given rise to the reallocation phenotype. This could happen by one of the two possible mechanisms. First, cold shock-induced damage may signal to the worm that death is imminent and reproductive timespan is limited. Worms may then funnel all their resources into embryos which are then better capable of surviving cold stress. However, this seems unlikely since these data suggest that *C. elegans* that avoid reallocation (or are genetically prevented from doing so) survive CS extremely well, which is inconsistent with CS causing large amounts of damage. It is possible that these worms are simply more resistant to CS-induced damage or are better able to repair damage in the absence of reallocation, though this would need to be tested more rigorously.

A second, more plausible scenario is that seasonal cold cycles signal a coming winter season with a low chance of offspring survival. At specific larval stages, worms can enter a highly stress-resistant state (called the dauer stage) which may enable them to survive the winter season and resume development when conditions are more favorable. If the better-provisioned embryos of worms that have recently experienced a cold cycle survive successive cold shock, this could favor a larger percentage of offspring that are able to successfully enter the dauer state. Long-term, this would translate to a higher probability that offspring survive to reproduce when winter conditions are alleviated, continuing the parent’s lineage. This model accounts for the selection of the reallocation trait in evolutionary history. It further suggests that rather than being induced as a result of damage that signals impending death (and thus a final chance to reproduce), reallocation may instead be prompted by a more general prediction of future damage to both the parent and the offspring. Further experimentation is needed to assess this model and to determine how it would fit into the known framework for terminal investment strategies.

Although we have largely considered simple maternal effects involving cytosolic investments in this work, there is also the potential for epigenetic effects at the level of gene expression to play a role in intergenerational and transgenerational offspring survival. Indeed, the progeny of *C. elegans* submitted to various stressors demonstrate improvements in stress survival to the same and other types of stress. Recently, a study by Burton et al. [7] examined multigenerational responses to a number of stressors including nutrient deprivation and pathogenic stress and found that multiple *Caenorhabditis* species regulate gene expression in the F1 generation in a stress-specific manner to benefit survival upon exposure to the same. These studies highlight the evolutionary advantage of provisioning offspring for what is perceived to be a likely future. It would be interesting to extend analyses of multigenerational gene regulation to see if this is also at play in the context of CS.

### Terminal investment has consequences for population-level dynamics

Terminal investment occurs across a broad phylogenetic range in response to many stressors, and the potential ecological and evolutionary implications are serious in today’s global climate. Pathogen exposure commonly elicits terminal investment; one particularly relevant example occurs in several species of frog in response to infection by *Batrachochytrium dendrobatidis* (Bd), the perpetrator of severe global amphibian declines. Rather than retaining resources for survival, individuals in many of these species direct energy to reproduction, increasing gamete output. While producing more offspring improves the likelihood of population persistence, this mode of reproduction does not favor survival of frogs past infection, whereupon animals selected by resilience would engender less susceptible offspring and gradually allow population resistance to the fungus to evolve [31]. Thus, terminal investment poses a serious long-term survival risk for some frog species faced with extinction from Bd. As global climates shift, it is unclear how population dynamics will be affected by changing temperatures. Since terminal investment is suspected to be a means of population persistence, it may play an important role in continued population survival of various species.

In most documented cases, terminal investment results in increases to brood size but some examples exist in which offspring quality is favored, such as in the tsetse fly, where stress levels are positively correlated with the percent of body fat that is dedicated to offspring [32]. The majority of terminal investment studies currently come from ecological studies in non-model systems; thus, an understanding of the cellular components and mechanisms involved in terminal investment, particularly in relation to quality investment, remains largely elusive. Here we have identified a new terminal investment process in response to cold stress in a genetically tractable and environmentally relevant model. The opportunity to better understand how this phenomenon is elicited and executed on the molecular scale is an exciting prospect for future studies. With a more comprehensive understanding of the impacts of stress on not only the parental generation but multiple generations thereafter, we can better predict both an organism’s physiology and population dynamics in response to specific environmental factors.

## Materials and Methods

### Strains and maintenance

All strains were maintained on Nematode Growth Medium (NGM) seeded with *Escherichia coli* OP50, at 20°C unless otherwise noted [33]. Worms were well-fed for at least three generations before any experiments. Some strains were provided by the CGC, which is funded by NIH Office of Research Infrastructure Programs (P40 OD010440). CGC strains used in this study were N2 (Bristol) wild-type, SJ4005 *zcIs4[hsp-4::gfp],* BR5514 *tax-2*(*p671*); *tax-4*(*p678*), GR2245 *skn-1*(*mg570*), RB2365 *vit-2*(*ok3211*), RB2382 *vit-5*(*ok3239*), CB4037 *glp-1*(*e2141*), and JK509 *glp-1*(*q231*). JRP1036 *skn-1*(*lax188*) was generated from SPC168 *dvIs19*; *skn-1*(*lax188*).

### Cold shock survival

For adult cold shock, approximately 20-70 young adult worms (not yet gravid) were picked to OP50-seeded 3.5 cm NGM and placed at 2°C for 4 hours. After 4 hours cold shock, plates were transferred to 20°C for worm recovery. Worms were scored at 1, 4, 24, 48, 72, and 96 hours post-cold shock for survival and qualitative phenotypic assessments. A worm was considered dead when nose tap did not elicit any movement. Clear worms were determined by a lack of almost all intestinal pigmentation, and immobile worms by a nose tap that elicited only movement in the head region.

For embryonic cold shock, parents were cold shocked as above, allowed to recover for 15 hours, then transferred to fresh plates to lay embryos for 2 hours. This time window is the peak of lipid reallocation following cold shock. Approximately 50-100 embryos were transferred to very lightly seeded 3.5 cm NGM plates and cold shocked at 2°C for 24 hours. Following a 24-hour recovery at 20°C, the number of hatched and unhatched embryos were counted. Embryos were considered hatched if the entirety of the L1 larva was visible in a non-curled state, and the larva was not dead.

During all cold shock experiments, plates were placed directly on the incubator shelf in a monolayer rather than a stack both during cold shock and the early stages of recovery to facilitate uniform temperature changes among plates.

### C. elegans Fluorescence Imaging and Quantification

zcIs4[*hsp-4::GFP*] worms were either heat shocked at 35°C for 2 h, cold shocked according to standard protocol, or non-shocked. After 12 h recovery at 20°C, worms were picked to a drop of M9 containing 5 mM sodium azide. Worms were imaged on a Nikon Eclipse 90i microscope at exposure times of 2.9 ms for DIC and 900 ms for GFP.

The fixed Nile Red staining protocol was modified from Pino et al. [34]. Briefly, at 12 h post-cold shock, 50-75 worms were washed once with M9 and fixed for three minutes at room temperature in 25 μl M9 + 150 μl 40% isopropanol. Worms were stained for 2 h in the dark with gentle rocking in 175 μl 40% isopropanol containing 0.6% Nile Red stock (0.5 mg/ml in acetone). Samples were washed with M9 for 30 min in the dark with gentle rocking, then mounted on a 2% agarose pad for imaging. Worms were imaged on a Nikon Eclipse 90i microscope at exposure times of 3 ms for DIC and 400 ms for GFP (figures 1, 3, 4, 5) or on a Zeiss Axio Imager Z1 microscope with exposure times of 3 ms for DIC and 500 ms for GFP (figures 2, 6).

Scale bars (100 μm) were added and all images were rotated and cropped using ImageJ. No other image manipulations were performed.

Images of fluorescent worms taken on the Nikon Eclipse 90i microscope (figures 1, 3, 4, 5) were analyzed using NIS Elements Data Analysis software; images taken on the Zeiss Axio Imager Z1 microscope (figures 2, 6) were analyzed using ImageJ. In either case, total fluorescence was determined by outlining the entire worm as a Region of Interest (ROI) and calculating the average fluorescence for that ROI. The number of internal embryos and the number of fluorescent embryos were determined by visual counts.

### Statistical analysis

All data represent at least three independent replicates. Statistical analysis was performed in GraphPad Prism 9 with an alpha value of P < 0.05. For samples with 2 conditions, nonparametric Mann-Whitney U tests were performed and reported with two-tailed P values. In cases with greater than 2 conditions, Kruskal-Wallis nonparametric ANOVA were performed with Dunn’s Multiple Comparison test. For hatching analyses, a Grubb’s test was conducted through GraphPad to identify outliers in the data, with one point identified as an outlier (Z = 3.4005, two-tailed P < 0.01) and removed. d’Agostino & Pearson normality tests were then applied to the remaining values and normal distribution was confirmed by a P > 0.05 for each condition. After validating data normality, 2-way ANOVAs were used to compare the mean of each condition with a Tukey’s Multiple Comparison test.

## Acknowledgements

The authors thank Sean Curran, Ralph Baumeister, and Thomas Heimbucher for reagents and useful discussions. Some strains were provided by the CGC, which is funded by NIH Office of Research Infrastructure Programs (P40 OD010440). Funding was provided by Gettysburg College.

## Author Contributions

LG and JRP conceived and performed experiments, analyzed and interpreted data, and prepared the manuscript.

